# microRNA-206 is a reproducibly sensitive and specific plasma biomarker of amyotrophic lateral sclerosis

**DOI:** 10.1101/2025.06.27.662023

**Authors:** Benjamin W. Henderson, Brian S. Roberts, Sherry Kolodziejczak, Meagan Cochran, Richard M. Myers

**Affiliations:** HudsonAlpha Institute for Biotechnology, 601 Genome Way, 35806, Huntsville, AL, USA; Crestwood Medical Center ALS Care Clinic, 610 Airport Road Suite 100, 35801, Huntsville, AL, USA

## Abstract

Amyotrophic lateral sclerosis (ALS) is a devastating and fatal neurodegenerative disease with no current therapeutic to modify disease progression. Reliable biomarkers for ALS are essential for improving diagnosis and evaluating therapeutic efficacy. We combined small-RNA sequencing from a discovery cohort of ALS patients and healthy controls with sequencing data from a previously published ALS cohort to identify candidate biomarkers. Machine learning analysis identified hsa-miR-206 as a strong classifier of ALS status in both cohorts. This finding was validated in an independent ALS cohort using droplet digital PCR (ddPCR), confirming the biomarker’s sensitivity and specificity in identifying ALS. Importantly, hsa-miR-206 also displayed high accuracy in differentiating ALS from Parkinson’s disease. These results further validate hsa-miR-206 as a circulating small-RNA biomarker for ALS with potential utility in diagnosis and therapeutic monitoring. Further studies in larger, diverse cohorts will be needed to validate its clinical applicability.

## Main

Amyotrophic lateral sclerosis (ALS) is a progressive neurodegenerative disease that affects approximately 200,000 individuals worldwide. Currently, therapeutic strategies focus on symptomatic treatment and do not affect disease progression, underscoring the urgent need for effective biomarkers and development of therapies^1,2^. Current biomarker approaches include imaging, neurophysiological studies, and fluid-based assays^2,3^. While imaging and neurophysiological-based biomarkers provide insight into structural and cell dysfunction aspects of ALS, respectively, they are costly, making repeated measurements and broad application impractical. Fluid biomarkers from cerebrospinal fluid, blood, and urine, on the other hand, offer a cost-effective and less invasive solution^2^. Much previous work in fluid-based biomarkers of ALS has been centered on the identification of proteins in circulation^4^. Neurofilament light (NfL) is an established biomarker of neuronal injury and is associated with disease severity and progression in ALS^5–8^. Other fluid biomarkers of ALS include those that measure alterations in immune regulation^9^, and the detection of gene products in circulation^10^. Often these biomarkers measure expression of a gene that is also a target of an experimental therapeutic. A biomarker that measures such a gene will thus report on target engagement efficacy without necessarily tracking disease alteration, limiting its usefulness.

Cell-free (cf) nucleic acids in plasma represent a promising alternative biomarker class. The abundance and composition of cfDNA or cfRNA often correlate with underlying physiological conditions and have increasingly become a target for “liquid biopsy” biomarker detection over the last decade^11–13^. In cancer research, for instance, cfDNA-based liquid biopsies are widely used to detect somatic mutations and monitor tumor dynamics^14,15^. Although these techniques offer high specificity and sensitivity, their utility is largely restricted to diseases where somatic mutations drive pathology, which does not apply to most ALS cases. More promising for ALS are cell-free RNA species, particularly small-RNAs (15–50 bp), which remain stable in the blood due to protection by proteins or encapsulation in exosomes^16^. Among small-RNAs, microRNAs (miRNAs) are especially attractive as biomarkers due to their role in post-transcriptional gene regulation and stability in circulation^17–19^. miRNAs, bound to the RNA-induced silencing complex (RISC) or encapsulated in extracellular vesicles, are detectable in plasma using sequencing or PCR-based methods^20^.

To agnostically and comprehensively identify small-RNA biomarkers, we assessed via sequencing all plasma-derived small-RNA classes as potential biomarkers of ALS status. As a discovery cohort, we collected plasma from patients enrolled at the Crestwood Medical Center ALS Care Clinic in Huntsville, AL, USA, under WGC IRB Protocol #1302717. The workflow for this study is outlined in Figure 1A. We enrolled a total of 99 patients, with 69 designated as healthy controls and 30 with clinician diagnosed ALS. Patient characteristics for both the healthy control and ALS cohorts can be found in Supplementary Table 1. We extracted total RNA from patient plasma, and prepared sequencing libraries as previously described^20^, with modifications outlined in Methods. We sequenced the libraries on a NextSeq2000, and processed FASTQs to count tables using a custom pipeline built with bash and Python scripts (see Supplementary Code). Sequencing generated an average of ∼10 million small-RNA aligned reads per sample (Supplementary Figure 1A), with a broad diversity of small-RNA species identified (Supplementary Figure 1A) that includes miRNAs, t-RNA derived fragments (tRfs), and other previously defined plasma-associated small-RNA species^20^. As part of the library preparation process, we blocked the inclusion of highly-abundant hsa-miR-451a and hsa-miR-16-5p using previously published methods^21^. These miRNAs derive from blood cell lysis and their reduction prevents their domination of sequencing reads, akin to ribosomal RNA depletion in mRNA libraries (Supplementary Figure 1B).

**Figure 1.**
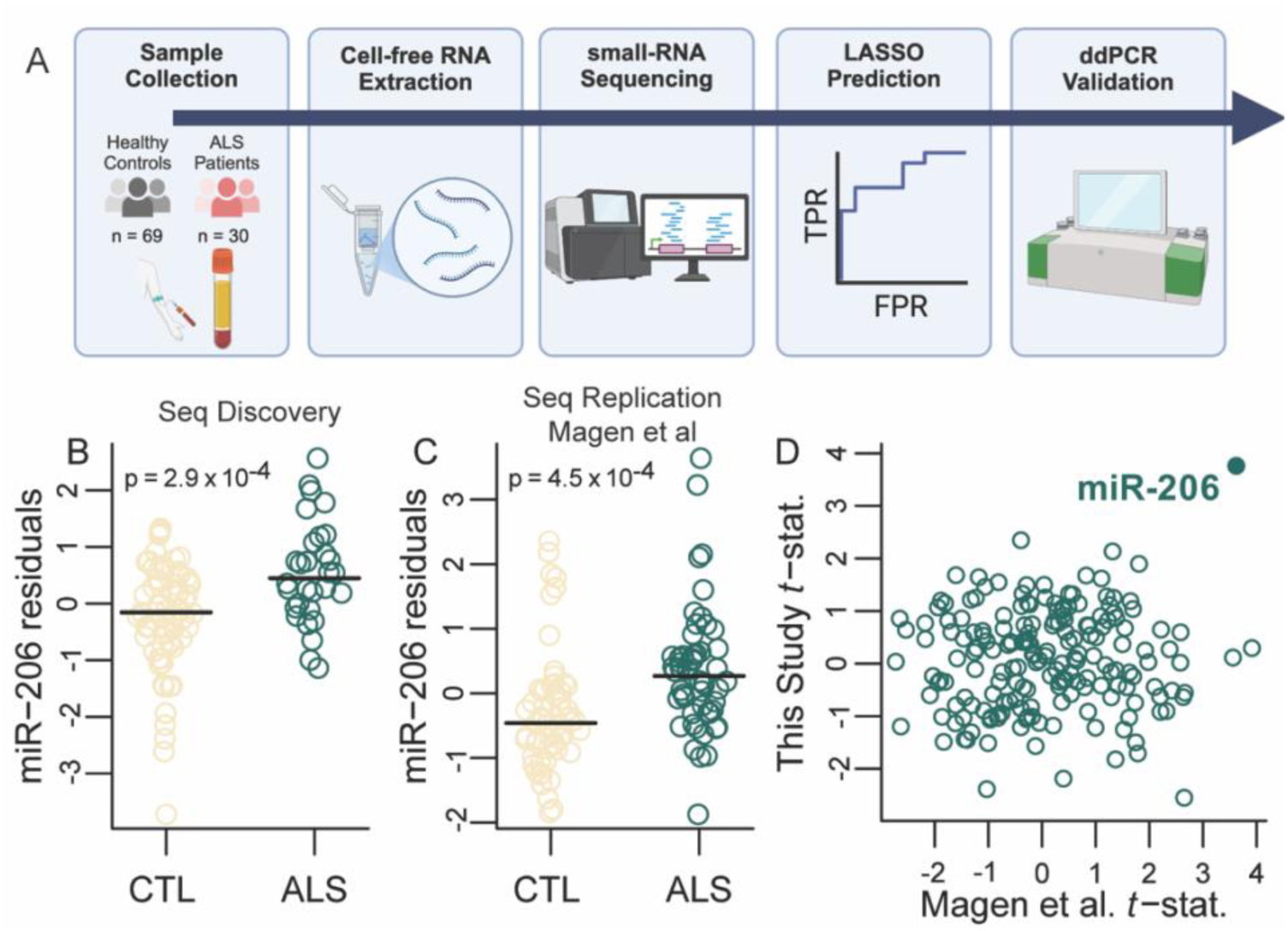
Identification of plasma miR-206 as a reproducible biomarker of ALS from sequencing studies. A) Graphical illustration of discovery cohort project workflow. In short, samples were processed through a high-throughput NGS workflow optimized to reduce bias in biomarker discovery, and validation was performed using ddPCR. B) Residuals from a linear regression of hsa-miR-206 log(counts per million) versus sample covariates (see Methods) are plotted for controls (“CTL”) and ALS from this study. P-value from t-test of regression coefficient (null model coefficient equals zero) is shown. C) Same as B but for data from Magen et al. (see Methods). D) For analytes passing filters in both sets (median counts greater than 50 and at least one count in every sample, n = 187) the *t*-statistic is plotted for each set.

To enhance our ability to identify differentially abundant small-RNAs between ALS patients and controls, we used count data from both this study and another publicly available dataset that we included as a secondary sequencing replication cohort^22^ (Magen *et al*.; GEO: GSE168714). We discarded libraries with fewer than 3 million aligned reads (this study) or 0.5 million (Magen *et al*.) and excluded libraries with high levels of the hemolysis marker hsa-miR-486-3p. Samples from batches lacking representation across sex and disease status were also removed. We retained only small-RNAs with a median read count >50 for reliability in follow-up ddPCR studies. Read depth and blood-cell lysis counts for the Magen *et al*. dataset can be found in Supplementary Figure 2. Batch effects were corrected using ComBat^23^, with covariates including disease status, sex, and age (Supplementary Tables 2 and 3). We fit linear models to each small RNA using sex, age, total aligned reads, and miR-486-3p CPM as independent variables. We then applied regularized elastic net logistic regression via R package glmnet^24^ on the residuals to identify optimal classifiers of ALS status (Supplementary Table 4). In both our sequencing discovery cohort and the Magen *et al*. sequencing replication cohort, hsa-miR-206 was identified as an optimal classifier of ALS disease status (Figure 1B-D). hsa-miR-206, known as a “myomiR”, is almost exclusively expressed in muscle tissue. Previous, less-powered studies associated higher abundance of miR-206 in plasma with ALS status^25–28^, consistent with our findings here. While we were able to associate other small-RNAs as significantly associated with ALS (Figure 1D, Supplementary Table 4), our analysis from two independent studies demonstrated that the highest performing predictive model used hsa-miR-206 alone as a classifier. We therefore focused only on hsa-miR-206 in further validation efforts.

To establish its veracity and clinical relevance, the validation of this sequencing-based signature required an orthogonal approach that does not share similar error mechanisms to sequencing; moreover, sequencing could prove difficult and costly to translate to clinical practice. Accordingly, we applied droplet digital PCR (ddPCR) to establish the feasibility of the clinical measurement of hsa-miR-206 as a biomarker of ALS. Other studies have focused on plasma small-RNA signature validation and clinical application using established approaches like RT-qPCR, as we have done^20,29^. Based on technological improvements and our own experience, ddPCR provides higher precision and a better lower limit of detection. We first applied ddPCR to a subset of healthy controls and ALS patients from our discovery (sequencing) cohort. Using the same RNA preparations that were used in sequencing, we prepared cDNA from each sample, generated droplets, and quantified hsa-miR-206 abundances. We z-scored (see Supplementary Code) absolute concentrations of hsa-miR-206 and observed higher quantities in ALS samples of hsa-miR-206 in agreement with the sequencing results (Figure 2B). Expanding upon the ability of hsa-miR-206 to correctly distinguish ALS from healthy controls, we trained a simple linear model on z-scored ddPCR expression data from our discovery cohort (Figure 2A). Our data show that hsa-miR-206 displays solid performance at distinguishing ALS cases from controls (AUC = 0.83, Figure 2A). We defined an optimal z-score threshold in which our model displays perfect specificity with good sensitivity (Figure 2A, red dot), and used this threshold moving forward when distinguishing ALS from healthy controls. Patient sex had no significant effect on hsa-miR-206 levels according to a simple regression analysis, and inclusion of sex into our model did not affect performance (Supplementary Figure 3).

**Figure 2.**
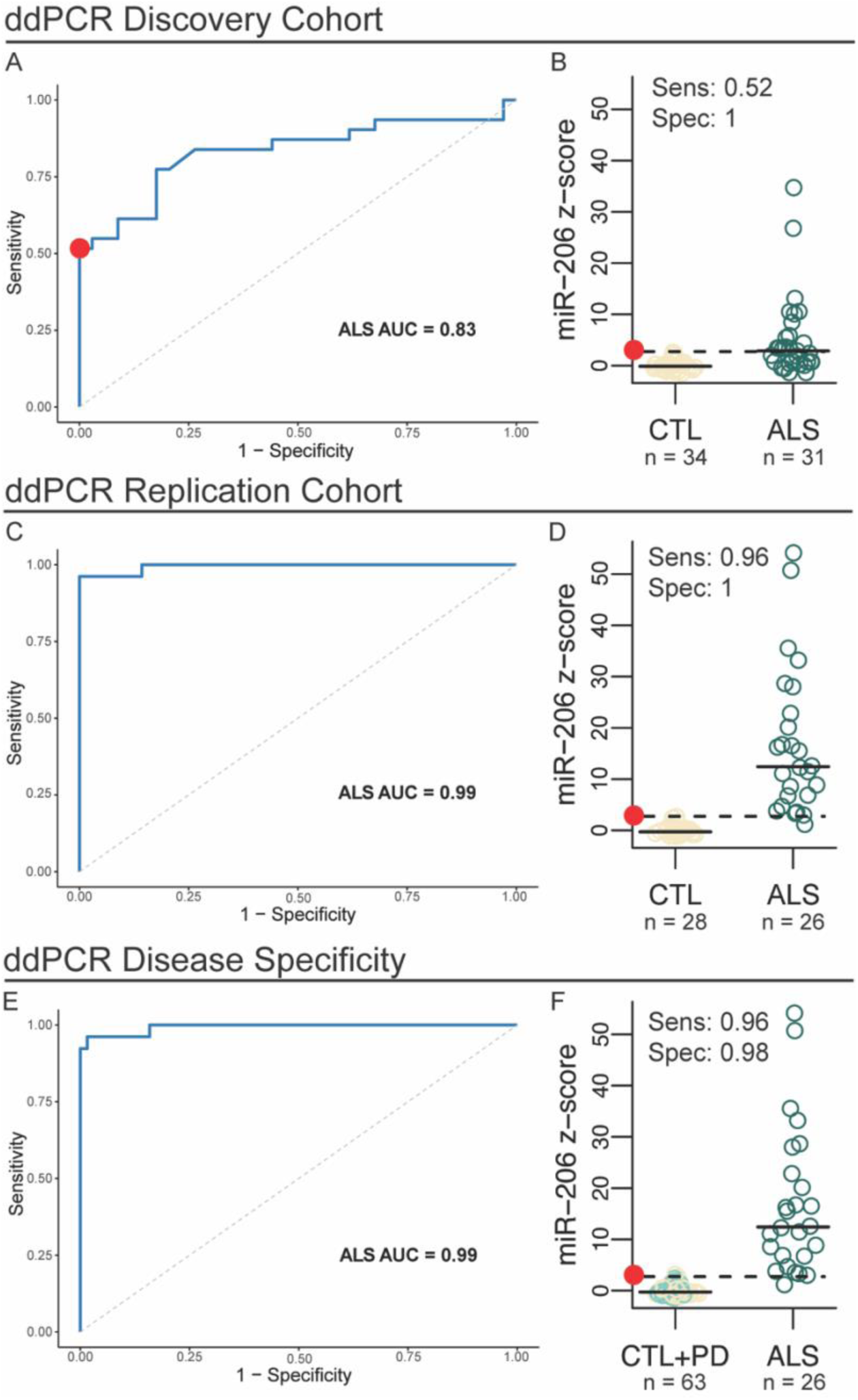
ddPCR analysis validates the ability of hsa-miR-206 to differentiate ALS from healthy controls and Parkinson’s disease. A) ROC curve displaying the ability of hsa-miR-206 to distinguish ALS from healthy controls in the discovery cohort based on ddPCR absolute quantification. The red dot represents the optimal threshold of sensitivity and specificity in the discovery cohort. B) Z-scored hsa-miR-206 ddPCR expression data, dashed lines represent the optimal threshold as shown in panel A. C-D) Same as A-B, but in the NEALS replication cohort. E-F) Same as A-B, but with the SFC Parkinson’s cohort included.

To further validate the ability of hsa-miR-206 to distinguish ALS patients from healthy controls, we acquired an independent replication cohort of samples from the Northeast Amyotrophic Lateral Sclerosis (NEALS) Biorepository (Supplementary Table 5). The cohort was age and sex-matched, and all ALS patients had a confirmed diagnosis from a physician. We extracted RNA from 500µL plasma and then prepared cDNA and performed ddPCR as previously described. Patient characteristics such as sex, collection site, and age at collection had no significant effect on hsa-miR-206 expression in our expanded analysis (Supplementary Figure 4). We found that hsa-miR-206 levels in ALS patients from the NEALS cohort followed the same trend as our sequencing data and were significantly elevated in ALS compared to healthy controls (Figure 2D). Of note, hsa-miR-206 levels as measured by ddPCR were overall higher in the NEALS cohort in both controls and patients. We hypothesize that this difference is likely due to blood collection tube differences in preserving levels of small-RNAs. We applied hsa-miR-206 ddPCR abundance as a binary classifier of ALS status using the threshold established in the Crestwood cohort to the NEALS cohort and ROC analysis revealed exceptional performance, with high sensitivity and specificity (0.96 and 1, respectively. Figure 2C-D).

A desirable trait of a biomarker is its specificity to the intended disease. To test the disease specificity of hsa-miR-206, we utilized a cohort of samples that were collected at the Smith Family Clinic at the HudsonAlpha Institute for Biotechnology that consisted of patients diagnosed with Parkinson’s disease and healthy controls (Supplementary Table 6). The cohort was age and sex-matched, and all PD patients had a confirmed diagnosis from a physician. We extracted RNA from 3 mL plasma and then prepared cDNA and performed ddPCR as previously described in this manuscript. We combined the z-scored hsa-miR-206 expression data for the Parkinson’s cohort with our NEALS cohort data and applied our model to this dataset. We observed that hsa-miR-206 still displayed exceptional performance in distinguishing ALS from controls, even with the addition of another neurodegenerative disease in the cohort, labeled for analysis as controls (Figure 2E, AUC = 0.99). Furthermore, when applying the optimal threshold for z-scored hsa-miR-206 expression, we observed high accuracy when distinguishing Parkinson’s cases from ALS (Figure 2F, accuracy = 0.97).

Considering the current state of ALS biomarkers, the utilization of small-RNA biomarkers in conjunction with the currently used protein and imaging-based biomarkers could provide a valuable complement to the field. Biomarkers that are not direct targets of therapeutics and report on cell-based reactions to therapies could potentially monitor therapeutic response that translates into altered disease progression. Cell-free small-RNAs provide a powerful approach for diagnosis, early detection and monitoring progress in response to therapeutics. In conclusion, our sequencing and ddPCR analyses have demonstrated high utility of hsa-miR-206 as a blood-based biomarker to distinguish ALS from healthy controls and other neurodegenerative diseases. Further studies will continue to evaluate the ability of hsa-miR-206 to distinguish ALS disease state, and to distinguish ALS from disease mimics.

## Methods

### Patient enrollment and sample acquisition

#### Discovery cohort patient enrollment

Study subjects were enrolled at the Crestwood Medical Center ALS Care Clinic in Huntsville, AL, USA. All ALS patients were required to have a physician-confirmed diagnosis of ALS. Written informed consent was obtained from each patient in accordance with the approved protocol WGC IRB #1302717. This cohort was comprised of 69 patients designated as healthy controls and 30 patients with a clinical diagnosis of ALS. A total of approximately 16 mL of blood was collected from each patient, split between one Streck RNA Complete BCT and one Streck cell-free DNA BCT (Streck, La Vista, NE). Within 24 hrs of collection, samples were double-spun according to manufacturer’s instructions. The plasma supernatant was isolated, yielding approximately 3-5 mL of plasma that was immediately frozen at -80 C until RNA isolation. For each patient, detailed medical history and demographic information was obtained in accordance with WGC IRB #1302717, and can be found in Supplementary Table 1.

#### Replication cohort sample acquisition from NEALS

A second, independent cohort of patient plasma samples was acquired from the Northeast Amyotrophic Lateral Sclerosis Consortium (NEALS). This cohort included 26 patients diagnosed with ALS and 28 patents that were age- and sex-matched healthy controls. For each sample, 500 µL of plasma was received along with detailed demographic and patient data (see Supplementary Table 5).

#### Parkinson’s disease cohort

For the validation of the specificity of our biomarker to ALS, we acquired samples from a third cohort of patients that were sex- and age-matched and were diagnosed with Parkinson’s disease (n = 20) or classified as a healthy control (n = 15). These samples were collected at the Smith Family Clinic for Genomic Medicine at the HudsonAlpha Institute for Biotechnology in Huntsville, AL, USA. Detailed demographic information for this cohort can be found in Supplementary Table 6. These samples were collected under approved WGC IRB #1361026 and followed the same plasma processing protocol as outlined for the ALS samples collected as part of the discovery cohort.

#### RNA isolation and sequencing library preparation

Prior to RNA extraction, all samples were thawed on ice. Manufacturer’s recommendations were then followed, and 3 mL of plasma were then used as input into the Norgen Plasma/Serum Circulating and Exosomal RNA Purification Kit Slurry Format (Norgen Biotek) and the concentrated using the Norgen RNA Clean and Concentration Kit (Norgen Biotek). Manufacturer instructions were followed exactly for both kits, and RNA was stored at -80 C until further use. If plasma samples originated from a standard EDTA collection tube, then only 500 µLs of plasma were used as input into RNA extractions.

Libraries for small-RNA sequencing were prepared as previously described^20^ with minor modifications. Namely, four degenerate bases were included in the 3’ adaptor sequence to mitigate adaptor ligation bias. In short, each sample underwent adaptor ligation, cDNA generation, and 16 cycles of PCR amplification. In addition to these steps, we selectively targeted the microRNAs hsa-miR-16-5p and hsa-miR-451a using our previously described methods^21^ to ensure enhanced complexity of the small-RNA sequencing library. The end PCR product was then separated under extremely denaturing gel electrophoresis, and fragments were excised that corresponded to an insert size of 15-30 base pairs. Post-purification, library concentrations were determined using the KAPA Library Quantification Kit (KAPA Biosystems), and a 3 nM pool was submitted for sequencing. Libraries were sequenced by SeqCenter on a NextSeq2000 (Illumina) P3 or P4 flowcell. Library preparations were split into three separate, balanced groups to minimize batch effect. All raw fastq files and a count table for all samples have been posted to GEO.

#### Processing of small RNA sequence data

To process raw sequencing data from FASTQ files to count tables, we employed a custom pipeline that uses bash scripting and python (Supplementary Code). We removed adaptor sequence using cutadapt^30^. Next, we used Bowtie2^31^ to align the reads to a custom small RNA reference (Supplementary Code). We created this reference from miRbase version 22^32^, the tRNA fragment database tRFdb^33^, and from sequences with a mean count of ten in a large sample set from our previous publication^20^. A complete list of reference sequences is provided in “all_small_RNA_non_redundant_seqs.fa” in the Supplementary Code. We generated count tables by parsing aligned BAM files using python scripts.

#### Identification of differentially abundant small RNAs between ALS patients and controls

From counts tables from both this current study and from those generated by Magen et al. (GEO: GSE168714) we employed R scripts (R Core Team 2022) to find differentially abundant small RNAs between ALS patients and controls (Supplementary Code). We calculated counts per million (CPM) for each library. For this study, we eliminated libraries with less than 3 million total aligned reads (Supplementary Figure 1) and for Magen et al. with less than 0.5 million total aligned reads (Supplementary Figure 2). The Magen et al. libraries in general had far fewer aligned reads and a higher minimum total count threshold would have resulted in excessive sample loss. We also eliminated libraries with the hemolysis marker miR-486-3p^34^ levels greater than the 75th percentile for both studies (Supplementary Figures 1-2). We then removed any remaining libraries that were in a batch that did not have at least one of each class for disease status and sex. We restricted the small RNAs kept for further analysis to those with a median number of reads greater than 50. We have found in previous work that this level allows for reliable measurement with ddPCR in follow up studies.

We corrected the filtered data for batch effects using ComBat^23^ with disease status, sex, and age in a provided model matrix. For the Magen et al. data, we only used disease status and sex in the model matrix because age was not provided for control samples. We then applied linear regression via the lm R function to each batch-corrected small RNA values with a model comprising sex, age (not for Magen et al.), total library aligned reads, and miR-486-3p CPM. We used this approach because the downstream analysis using regularized logistic regression via LASSO does not accept covariates. Next, we ran a second linear regression on the resulting residuals with ALS disease status as sole independent variable, calculating t-statistics and p-values based on a null hypothesis that the regression coefficient equals zero. Finally, we ran regularized logistic regression via the R package glmnet^24^ to identify sets of small RNA measurements that comprise an optimal classifier of ALS disease status.

#### cDNA generation and droplet digital PCR

We used the Applied Biosystems TaqMan Advanced miRNA cDNA Synthesis Kit (#A28007) to generate cDNA prior to ddPCR. This kit uses a poly-adenylase and ligation of a 5’ adaptor to generate cDNA. For samples that were collected in Streck RNA Complete BCTs, 2 µLs of RNA were used as input into the 10 µL cDNA reaction. For samples collected in standard EDTA vacutainers, 3.7 µLs of RNA were used as input into the 10-µL reaction. The cDNA product was diluted 1:10 with molecular biology grade water prior to ddPCR. 9 µLs of diluted cDNA was combined with 10 µLs of Bio-Rad ddPCR Supermix for Probes (No dUTP), 1 µL of Applied Biosystems TaqMan Advanced miRNA Assay, and 2 µLs of molecular biology grade water to create a 22 µL reaction that was loaded into the Bio-Rad AutoDG Droplet Generator, according to manufacturer’s instructions. After droplet generation, samples immediately underwent 40 cycles of endpoint PCR, according to Bio-Rad ddPCR Supermix for Probes (No dUTP) manufacturer’s instructions. After PCR, droplets were read using the Bio-Rad QX200 droplet reader and the QuantaSoft software. Positivity thresholds were automatically determined by the software, as to reduce human error in analysis. Biological samples were run in technical duplicates, requiring an automated reading of one of the technical duplicates to be considered for further analysis.

#### Analysis of ddPCR data

From the raw ddPCR data generated in the QuantaSoft program, we generated an average concentration for each sample represented by copies/ µL of reaction. We then employed R scripts (R Core Team 2022) for normalization and further analysis of ddPCR expression data (Supplementary Code). Briefly, concentrations for each sample were z-scored within their own cohort. Simple linear regression was performed to determine if covariates such as sex, age, and collection site affected expression data. A logistic regression model was then fit using z-scored data as a predictor of ALS status. ROC curves were generated using the R package pROC. An optimal threshold was determined from the discovery cohort by identifying the value that obtained the highest specificity and was then applied as the optimal cutoff for distinguishing ALS from control. The optimal threshold was then applied to all replication cohorts to obtain the sensitivity, specificity, or accuracy of distinguishing ALS from either control of Parkinson’s disease patients.

---

**Supplementary Figure 1.**
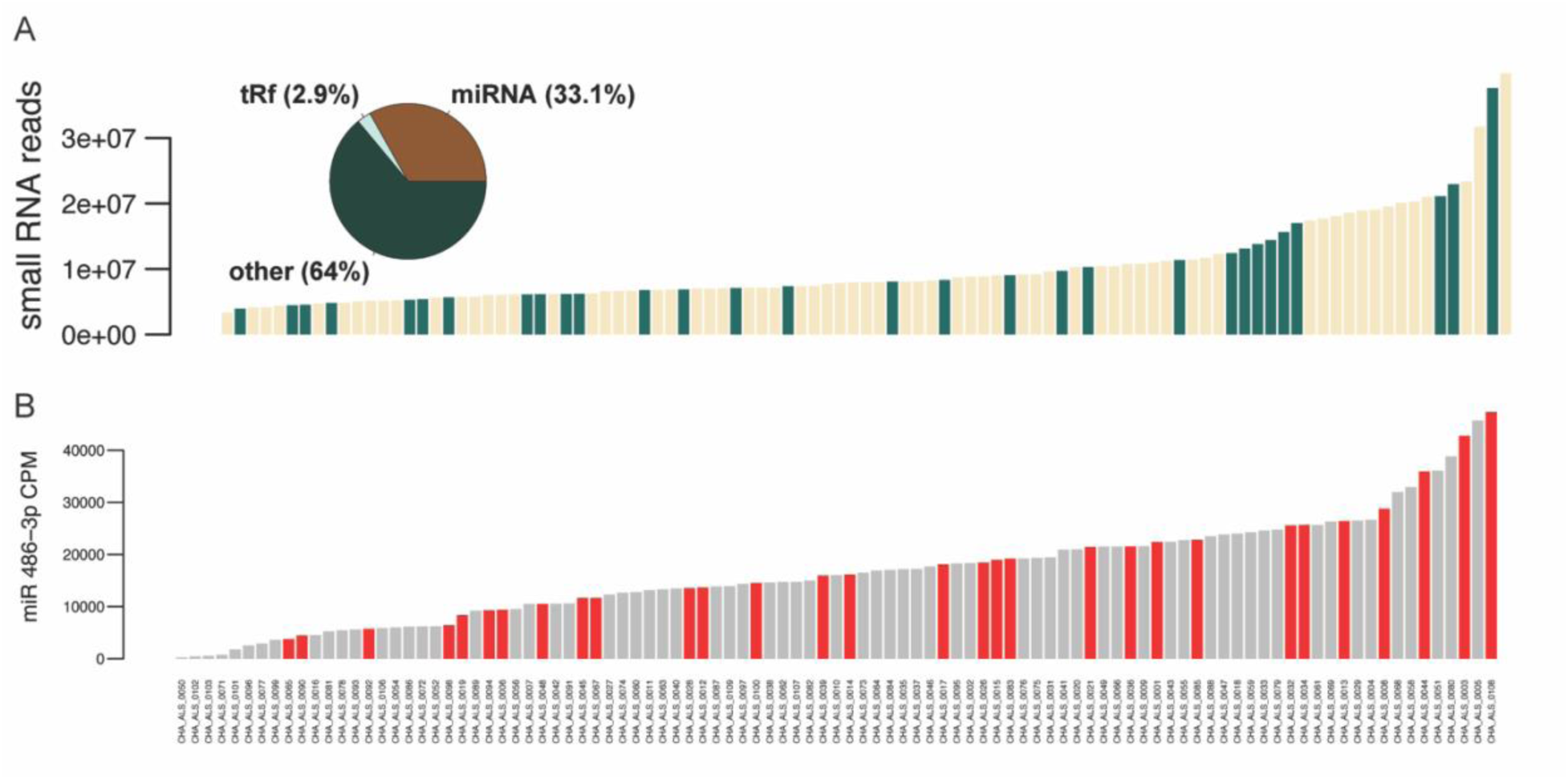
Quality control sequencing metrics. A) An average of ∼10 million small-RNA reads were aligned per sequencing library, with a diverse representation of miRNAs, tRfs, and “other” circulating small-RNA species. B) Quantities of a red blood cell miRNA marker in Crestwood-HudsonAlpha cohort sequencing libraries. Bar heights indicate counts per million (CPM) of hsa-mIR-486-3p, a marker of red blood cells and an indicator of the degree of cell lysis in the plasma sample. Gray bars indicate samples form controls and red bars indicate samples from ALS patients.

**Supplementary Figure 2.**
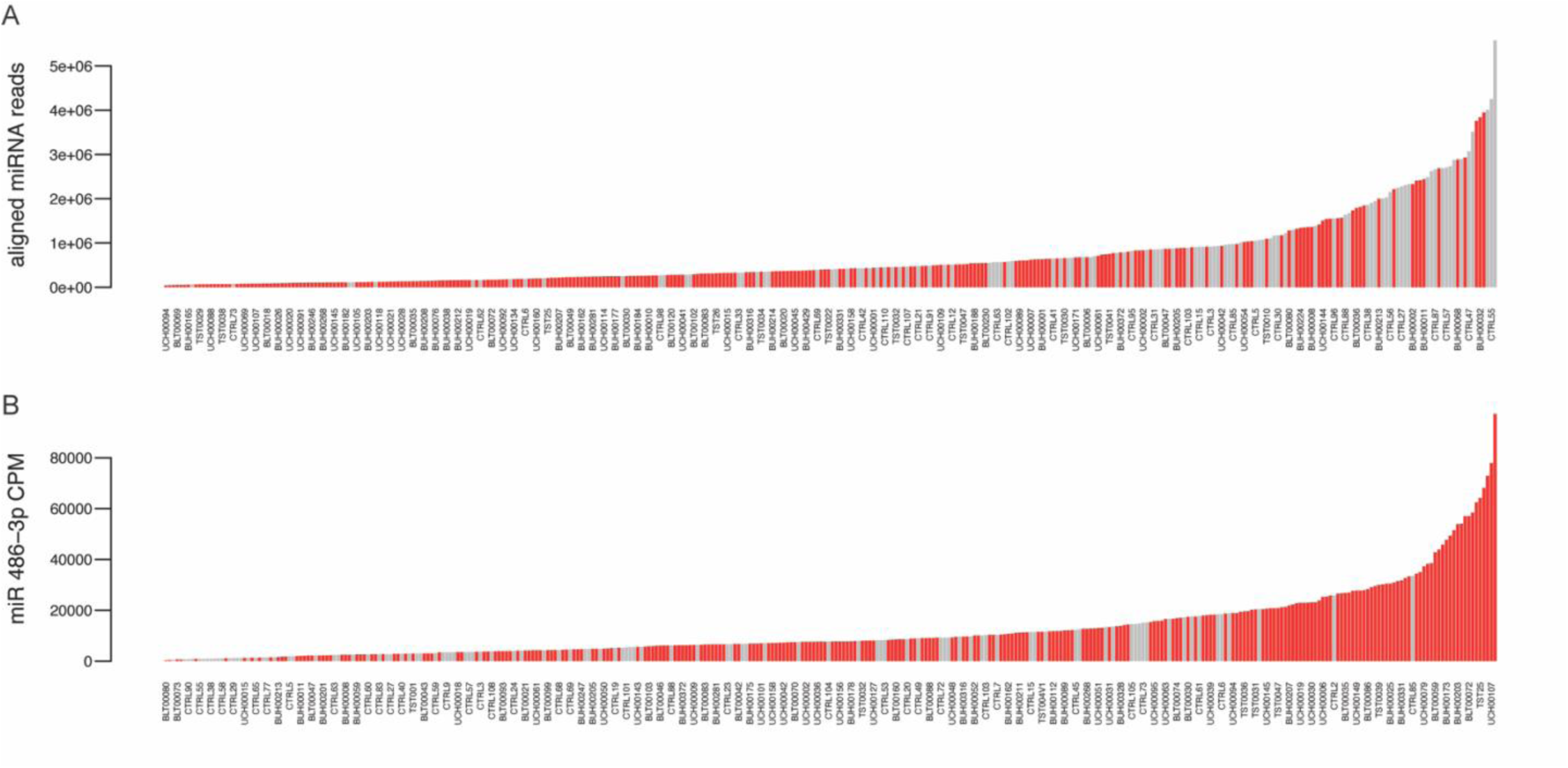
Quantities of total aligned miRNAs and a of red blood cell miRNA marker in Magen et al. cohort sequencing libraries. A) Bar heights indicate number of reads aligning to miRNAs for each sample library. B) Bar heights indicate counts per million (CPM) of hsa-mIR-486-3p, a marker of red blood cells and an indicator of the degree of cell lysis in the plasma sample. Gray bars indicate samples from controls and red bars indicate samples from ALS patients.

**Supplementary Figure 3.**
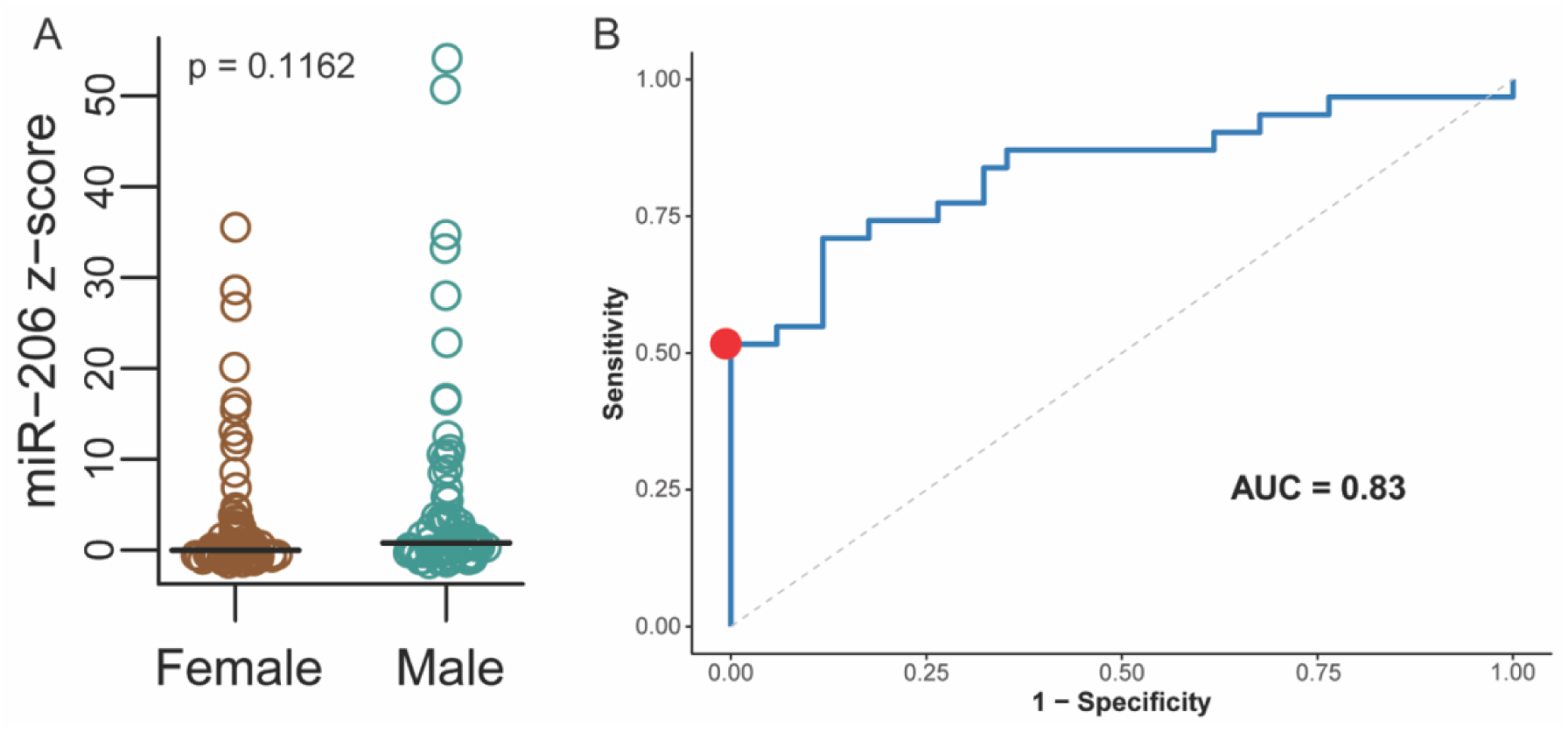
Sex does not have a significant effect on hsa-miR-206 expression or ability to differentiate disease from control. A) Z-scored hsa-miR-206 expression in all samples analyzed by ddPCR. The p-value was computed from a t-test of the regression coefficient, testing the association between hsa-miR-206 expression and sex. B) ROC curve displaying the ability of hsa-miR-206 to distinguish ALS from healthy controls in the discovery cohort based on ddPCR absolute quantification, with sex included in the analysis.

**Supplementary Figure 4.**
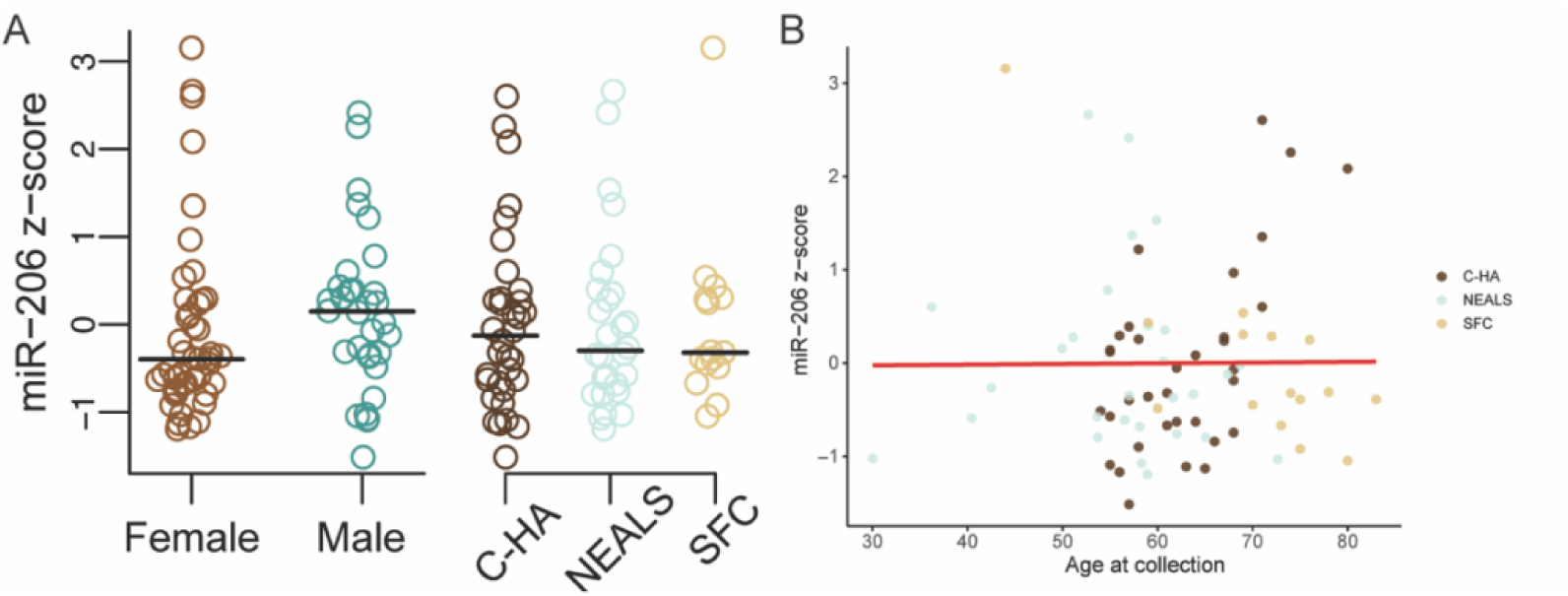
Association of control patient characteristics with hsa-miR-206 expression. A) hsa-miR-206 z-scored expression data for controls represented by sex and cohort. Linear regression analysis shows no significant association between hsa-miR-206 expression and sex or cohort. B) hsa-miR-206 z-scored expression data for controls plotted against age at collection. Linear regression analysis shows no significant association between hsa-miR-206 expression and age at collection.

**Supplementary Table 1.** Patient characteristiics from the Crestwood discovery cohort. Cohort characteristics from Healthy Control and ALS patients from the discovery cohort collected at the Crestwood Medical Center ALS Center of Excellence in Huntsville, AL.

|  | Healthy<br>Control | ALS |
| --- | --- | --- |
| Age; mean; y | 50.6 | 60.7 |
| Sex; n (%) |  |  |
| Female | 47 (68.12) | 12 (40) |
| Male | 22 (31.88) | 18 (60) |
| Race; n (%) |  |  |
| White | 69 (100) | 30 (100) |
| Disease Onset; n(%) |  |  |
| Limb |  | 13 (43.3) |
| Bulbar |  | 9(30) |
| Both |  | 2 (6.7) |
| Unknown |  | 6 (20) |

**Supplementary Table 2.** Association of sample characteristics with principal components in the Crestwood-HudsonAlpha cohort. Principal components were calculated on log(CPM) for all samples and detected small RNAs. A linear regression with all variables was run. The values are the p-values for each variable’s association with the given principal component. Red shading indicates a p-value less than 0.01.

|  | PC1 | PC2 | PC3 | PC4 | PC5 | PC6 | PC7 | PC8 | PC9 | PC10 |
| --- | --- | --- | --- | --- | --- | --- | --- | --- | --- | --- |
| **Batch** | 3.94E-05 | 4.60E-24 | 1.10E-05 | 8.27E-05 | 5.29E-01 | 6.76E-01 | 8.50E-01 | 4.30E-01 | 2.87E-01 | 8.84E-01 |
| **Sex** | 4.04E-01 | 8.98E-02 | 3.23E-01 | 5.50E-01 | 1.75E-02 | 2.78E-02 | 3.83E-01 | 6.49E-01 | 2.28E-01 | 5.00E-01 |
| **ALS Status** | 4.48E-01 | 5.08E-03 | 7.42E-01 | 6.94E-01 | 1.61E-01 | 4.02E-02 | 4.17E-01 | 2.90E-01 | 5.86E-01 | 1.33E-01 |
| **log(Total Counts)** | 2.62E-01 | 1.18E-01 | 4.08E-20 | 1.56E-01 | 9.86E-01 | 5.54E-01 | 4.85E-01 | 4.69E-02 | 8.83E-01 | 7.16E-01 |
| **log( miR-486 CPM)** | 2.49E-03 | 1.75E-06 | 4.98E-01 | 6.56E-12 | 5.27E-01 | 1.90E-01 | 1.31E-01 | 6.83E-01 | 1.30E-02 | 4.90E-01 |

**Supplementary Table 3.** Association of sample characteristics with principal components in the Magen et al. cohort. Principal components were calculated on log(CPM) for all samples and detected small RNAs. A linear regression with all variables was run. The values are the p-values for each variable’s association with the given principal component. Red shading indicates a p-value less than 0.01.

|  | PC1 | PC2 | PC3 | PC4 | PC5 | PC6 | PC7 | PC8 | PC9 | PC10 |
| --- | --- | --- | --- | --- | --- | --- | --- | --- | --- | --- |
| **Batch** | 2.34E-93 | 1.75E-23 | 2.34E-87 | 2.54E-63 | 1.40E-10 | 1.27E-05 | 3.50E-27 | 1.62E-01 | 7.32E-07 | 1.56E-01 |
| **Sex** | 3.55E-01 | 8.23E-02 | 5.91E-01 | 4.76E-01 | 4.89E-01 | 1.86E-01 | 4.85E-01 | 3.24E-01 | 5.78E-02 | 8.56E-01 |
| **ALS Status** | 6.20E-11 | 1.10E-03 | 5.81E-08 | 5.86E-04 | 1.27E-01 | 9.14E-01 | 9.49E-01 | 3.85E-01 | 6.19E-01 | 1.00E-01 |
| **Age of Onset** | 2.72E-01 | 4.15E-01 | 5.46E-01 | 4.51E-02 | 8.85E-01 | 8.07E-01 | 9.90E-01 | 6.86E-01 | 9.54E-02 | 7.00E-01 |
| **Riluzole Treatment** | 3.30E-01 | 4.58E-01 | 7.98E-02 | 7.25E-01 | 8.13E-01 | 9.83E-01 | 4.75E-01 | 1.33E-01 | 7.55E-01 | 9.67E-01 |
| **log(Total Counts)** | 2.48E-05 | 3.83E-44 | 1.01E-07 | 4.24E-01 | 1.02E-06 | 2.01E-02 | 3.52E-02 | 1.39E-02 | 4.78E-02 | 2.29E-02 |
| **log( miR-486 CPM)** | 7.63E-03 | 7.41E-43 | 4.88E-05 | 8.52E-01 | 1.60E-07 | 6.59E-01 | 1.47E-02 | 1.44E-02 | 5.36E-02 | 1.78E-02 |

**Supplementary Table 4.**
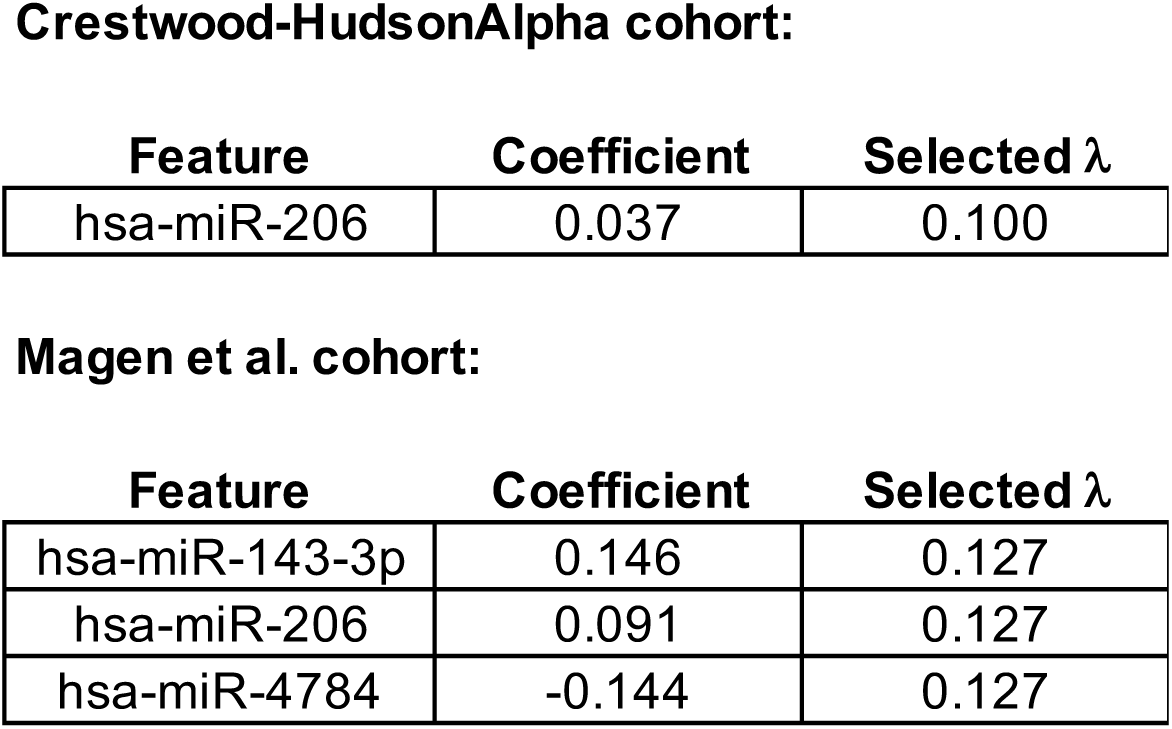
Selected features from LASSO regression. LASSO regression was applied via the R package “glmnet” to selected features from each cohort. The optimal set was selected from the lambda value of “lambda.1se”.

**Crestwood-HudsonAlpha cohort:**
| Feature | Coefficient | Selected $\lambda$ |
| --- | --- | --- |
| hsa-miR-206 | 0.037 | 0.100 |

| Feature | Coefficient | Selected $\lambda$ |
| --- | --- | --- |
| hsa-miR-143-3p | 0.146 | 0.127 |
| hsa-miR-206 | 0.091 | 0.127 |
| hsa-miR-4784 | -0.144 | 0.127 |

**Supplementary Table 5.**
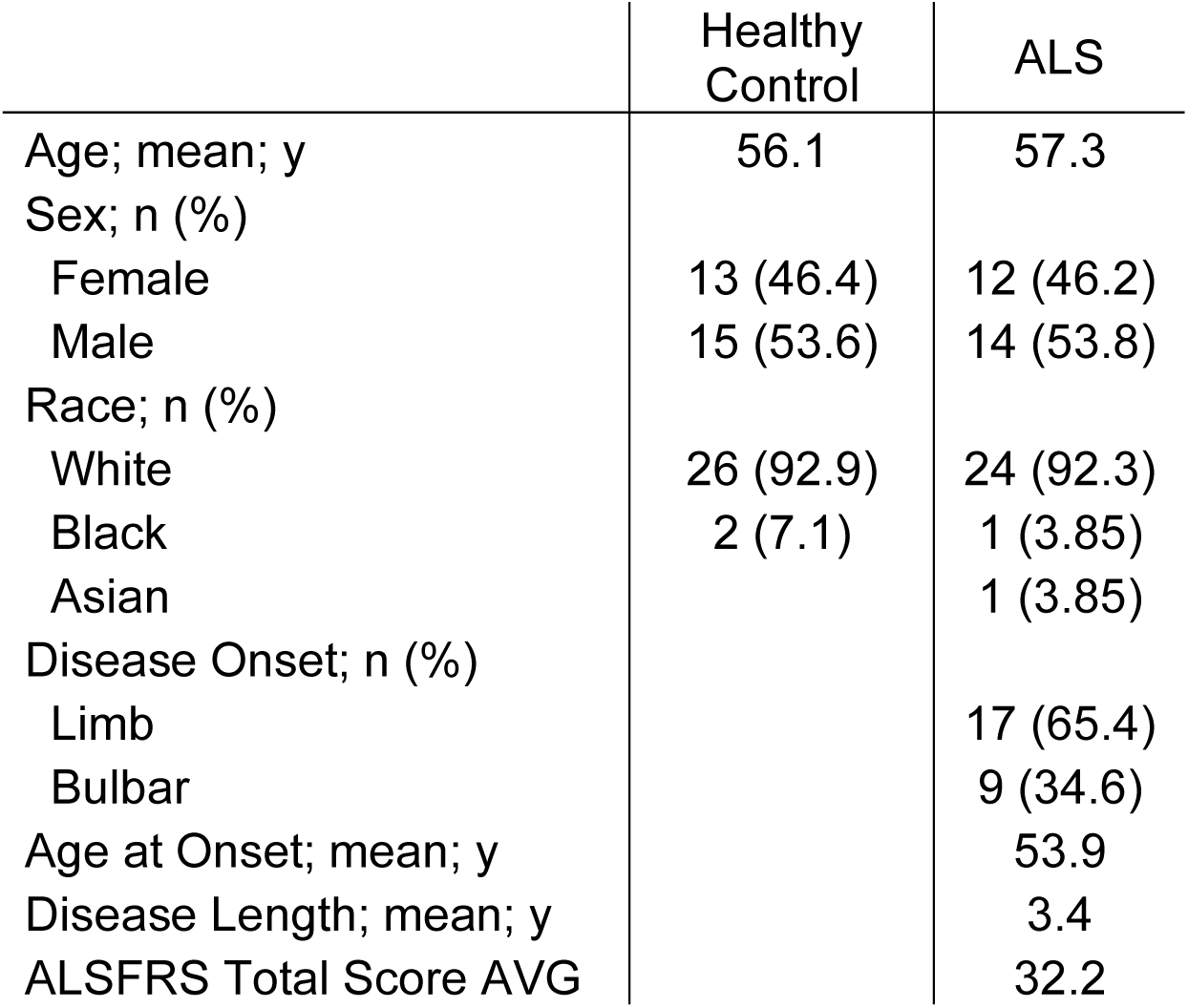
Patient characteristics from the NEALS replication cohort. Cohort characteristics from Healthy Control and ALS patients from the replication cohort acquired from the Northeast Amyotrophic Lateral Sclerosis (NEALS) Consortium.

|  | Healthy<br>Control | ALS |
| --- | --- | --- |
| Age; mean; y | 56.1 | 57.3 |
| Sex; n (%) |  |  |
| Female | 13 (46.4) | 12 (46.2) |
| Male | 15 (53.6) | 14 (53.8) |
| Race; n (%) |  |  |
| White | 26 (92.9) | 24 (92.3) |
| Black | 2 (7.1) | 1 (3.85) |
| Asian |  | 1 (3.85) |
| Disease Onset; n (%) |  |  |
| Limb |  | 17 (65.4) |
| Bulbar |  | 9 (34.6) |
| Age at Onset; mean; y |  | 53.9 |
| Disease Length; mean; y |  | 3.4 |
| ALSFRS Total Score AVG |  | 32.2 |

**Supplementary Table 6.**
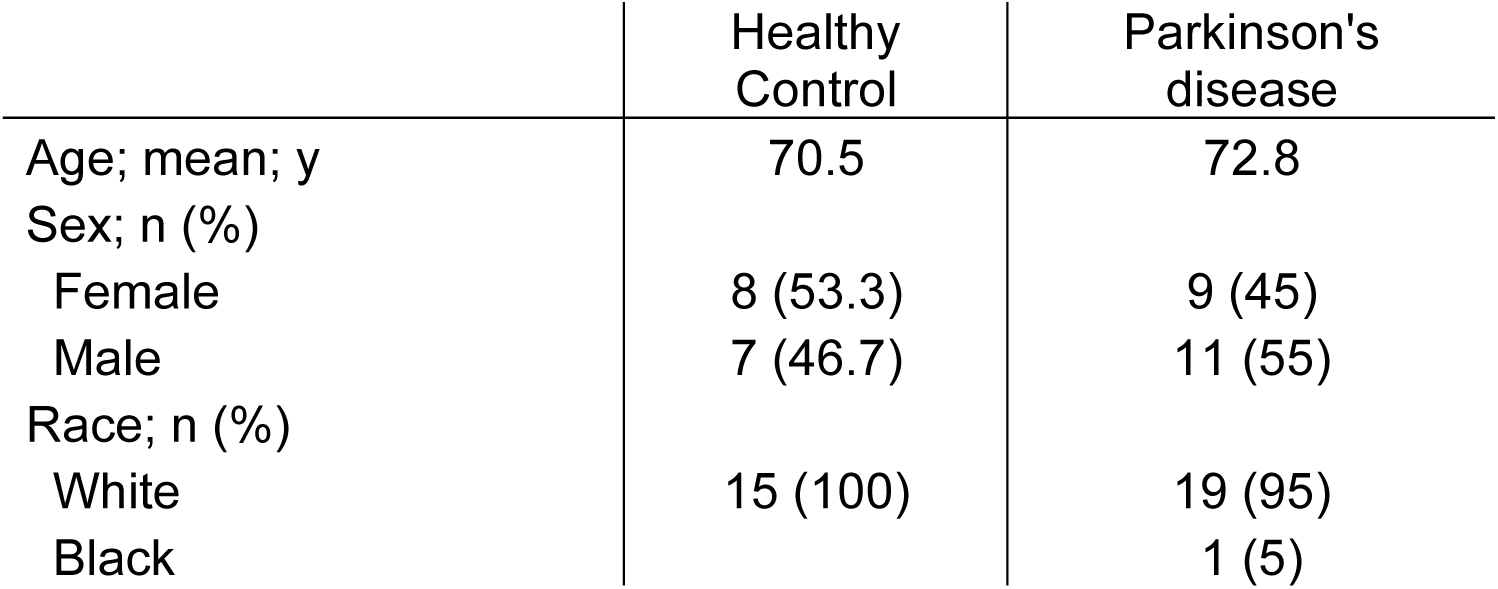
Patient characteristics from the Smith Family Clinic Parkinson’s disease cohort. Cohort characteristics from Healthy Control and Parkinson’s disease patients from the validation cohort collected at the Smith Family Clinic for Genomic Medicine at the HudsonAlpha Institute for Biotechnology.

|  | Healthy Control | Parkinson's disease |
| --- | --- | --- |
| Age; mean; y | 70.5 | 72.8 |
| Sex; n (%) |  |  |
| Female | 8 (53.3) | 9 (45) |
| Male | 7 (46.7) | 11 (55) |
| Race; n (%) |  |  |
| White | 15 (100) | 19 (95) |
| Black |  | 1 (5) |

## Literature Cited

1. Lu, L., Deng, Y. & Xu, R. Current potential therapeutics of amyotrophic lateral sclerosis. Front. Neurol. 15, (2024).

2. Irwin, K. E., Sheth, U., Wong, P. C. & Gendron, T. F. Fluid biomarkers for amyotrophic lateral sclerosis: a review. Mol Neurodegeneration 19, 9 (2024).

3. McMackin, R., Bede, P., Ingre, C., Malaspina, A. & Hardiman, O. Biomarkers in amyotrophic lateral sclerosis: current status and future prospects. Nat Rev Neurol 19, 754–768 (2023).

4. Schreiber, S. et al. Significance of CSF NfL and tau in ALS. J Neurol 265, 2633–2645 (2018).

5. Delaby, C. et al. Differential levels of Neurofilament Light protein in cerebrospinal fluid in patients with a wide range of neurodegenerative disorders. Sci Rep 10, 9161 (2020).

6. Gaiani, A. et al. Diagnostic and Prognostic Biomarkers in Amyotrophic Lateral Sclerosis. JAMA Neurol 74, 525–532 (2017).

7. Lu, C.-H. et al. Neurofilament light chain. Neurology 84, 2247–2257 (2015).

8. Dreger, M. et al. Cerebrospinal Fluid Neurofilament Light Chain (NfL) Predicts Disease Aggressiveness in Amyotrophic Lateral Sclerosis: An Application of the D50 Disease Progression Model. Front Neurosci 15, 651651 (2021).

9. McCauley, M. E. & Baloh, R. H. Inflammation in ALS/FTD pathogenesis. Acta Neuropathol 137, 715–730 (2019).

10. Irwin, K. E. et al. A fluid biomarker reveals loss of TDP-43 splicing repression in presymptomatic ALS–FTD. Nat Med 30, 382–393 (2024).

11. Gaitsch, H., Franklin, R. J. M. & Reich, D. S. Cell-free DNA-based liquid biopsies in neurology. Brain 146, 1758–1774 (2022).

12. Roskams-Hieter, B. et al. Plasma cell-free RNA profiling distinguishes cancers from pre-malignant conditions in solid and hematologic malignancies. *npj Precis*. Onc. 6, 1–11 (2022).

13. Dube, U. et al. An atlas of cortical circular RNA expression in Alzheimer disease brains demonstrates clinical and pathological associations. Nat Neurosci 22, 1903–1912 (2019).

14. Tarazona, N. et al. Targeted next-generation sequencing of circulating-tumor DNA for tracking minimal residual disease in localized colon cancer. Ann Oncol 30, 1804–1812 (2019).

15. Coombes, R. C. et al. Personalized Detection of Circulating Tumor DNA Antedates Breast Cancer Metastatic Recurrence. Clin Cancer Res 25, 4255–4263 (2019).

16. Pratt, A. J. & MacRae, I. J. The RNA-induced silencing complex: a versatile gene-silencing machine. J Biol Chem 284, 17897–17901 (2009).

17. Saliminejad, K., Khorram Khorshid, H. R., Soleymani Fard, S. & Ghaffari, S. H. An overview of microRNAs: Biology, functions, therapeutics, and analysis methods. J Cell Physiol 234, 5451– 5465 (2019).

18. Arroyo, J. D. et al. Argonaute2 complexes carry a population of circulating microRNAs independent of vesicles in human plasma. Proc Natl Acad Sci U S A 108, 5003–5008 (2011).

19. Yu, X., Odenthal, M. & Fries, J. W. U. Exosomes as miRNA Carriers: Formation-Function-Future. Int J Mol Sci 17, 2028 (2016).

20. Roberts, B. S. et al. Discovery and Validation of Circulating Biomarkers of Colorectal Adenoma by High-Depth Small RNA Sequencing. Clin Cancer Res 24, 2092–2099 (2018).

21. Roberts, B. S. et al. Blocking of targeted microRNAs from next-generation sequencing libraries. Nucleic Acids Res 43, e145 (2015).

22. Magen, I. et al. Circulating miR-181 is a prognostic biomarker for amyotrophic lateral sclerosis. Nat Neurosci 24, 1534–1541 (2021).

23. Zhang, Y., Parmigiani, G. & Johnson, W. E. ComBat-seq: batch effect adjustment for RNA-seq count data. NAR Genomics and Bioinformatics 2, lqaa078 (2020).

24. Friedman, J. H., Hastie, T. & Tibshirani, R. Regularization Paths for Generalized Linear Models via Coordinate Descent. Journal of Statistical Software 33, 1–22 (2010).

25. Dobrowolny, G. et al. A longitudinal study defined circulating microRNAs as reliable biomarkers for disease prognosis and progression in ALS human patients. Cell Death Discov. 7, 1–11 (2021).

26. Gomes, B. C. et al. Differential Expression of miRNAs in Amyotrophic Lateral Sclerosis Patients. Mol Neurobiol 60, 7104–7117 (2023).

27. Toivonen, J. M. et al. MicroRNA-206: A Potential Circulating Biomarker Candidate for Amyotrophic Lateral Sclerosis. PLoS One 9, e89065 (2014).

28. Waller, R. et al. Serum miRNAs miR-206, 143-3p and 374b-5p as potential biomarkers for amyotrophic lateral sclerosis (ALS). Neurobiol Aging 55, 123–131 (2017).

29. Vaklavas, C. et al. TBCRC 002: a phase II, randomized, open-label trial of preoperative letrozole with or without bevacizumab in postmenopausal women with newly diagnosed stage 2/3 hormone receptor-positive and HER2-negative breast cancer. Breast Cancer Res 22, 22 (2020).

30. Martin, M. Cutadapt removes adapter sequences from high-throughput sequencing reads. EMBnet.journal 17, 10–12 (2011).

31. Langmead, B. & Salzberg, S. L. Fast gapped-read alignment with Bowtie 2. Nat Methods 9, 357–359 (2012).

32. Kozomara, A., Birgaoanu, M. & Griffiths-Jones, S. miRBase: from microRNA sequences to function. Nucleic Acids Res 47, D155–D162 (2019).

33. Kumar, P., Mudunuri, S. B., Anaya, J. & Dutta, A. tRFdb: a database for transfer RNA fragments. Nucleic Acids Res 43, D141–D145 (2015).

34. Pritchard, C. C. et al. Blood cell origin of circulating microRNAs: a cautionary note for cancer biomarker studies. Cancer Prev Res (Phila) 5, 492–497 (2012).

